# A novel polyphasic identification system for genus Trichoderma

**DOI:** 10.1101/481382

**Authors:** Kai Dou, Zhixiang Lu, Qiong Wu, Mi Ni, Chuanjin Yu, Meng Wang, Yaqian Li, Xinhua Wang, Huilan Xie, Jie Chen

## Abstract

As the rapid-changing in *Trichoderma* taxonomy, an efficient identification of *Trichoderma* species is an urgent issue for *Trichoderma*-based research. In this study, based on the current taxonomy of *Trichoderma*, we constructed curated databases including 338 ITS sequences, 435 TEF1 sequences, 415 RPB2 sequences and 28 phenotypic characters. In addition, a polyphasic identification system (PIST) was developed. Within PIST, the resolution ability of molecular and phenotype characters could be combined for species identification in a step-by-step way. Compared with other identification systems, the involved *Trichoderma* species were extended from 88 to 252 species and 175 from 188 tested *Trichoderma* species could be identified within PIST. In most tested cases, three nucleotide markers and phenotypic characters showed improved identification performance. The TEF1 sequences have superior resolution than other characters.

**Importance:** The genus *Trichoderma* is important to human society with a wide application in industry, agriculture and environment bio-remediation. Thus, a quick and accurate identification of *Trichoderma* spp. is paramount since it is usually the first step to conduct a *Trichoderma*-based scientific research and is an obstruction especially for those researchers as nematologist, chemists, nutritionist and the like, lacking of taxonomic knowledge of fungi.

## Introduction

The genus *Trichoderma* as a large group of microorganisms worldwide contains kinds of opportunistic fungi with economically and ecologically importance to human society. *Trichoderma* spp. has already been used for a long time in industrial enzyme production (C. et al, 2004; de Azevedo et al, 2000; Toyama et al, 2002), as biocontrol agents in biofertilizer and biopesticide (Contreras-Cornejo et al, 2009; Harman et al, 2004) or as bio-remediation agents for heavy metal and xenobiotic contamination (Tripathi et al, 2013; Zhang et al, 2018b). In addition, *Trichoderma* spp. is also employed to be an expression system for the production of heterologous proteins (Zhang et al, 2018a) and improve the feed nutrition for domestic animals (AlZahal et al, 2017). As scientific and practical values have continuously been explored, the studies involving *Trichoderma* spp. have been weaving through a variety of research fields and showing sustainable growth (Fig 1 and 2). Thus, accurate identification of *Trichoderma* spp. is paramount since it is usually the first step to conduct a scientific research and is an obstruction especially for those researchers as nematologist, chemists, nutritionist and the like, lacking of taxonomic knowledge of fungi.

**Fig 1.**
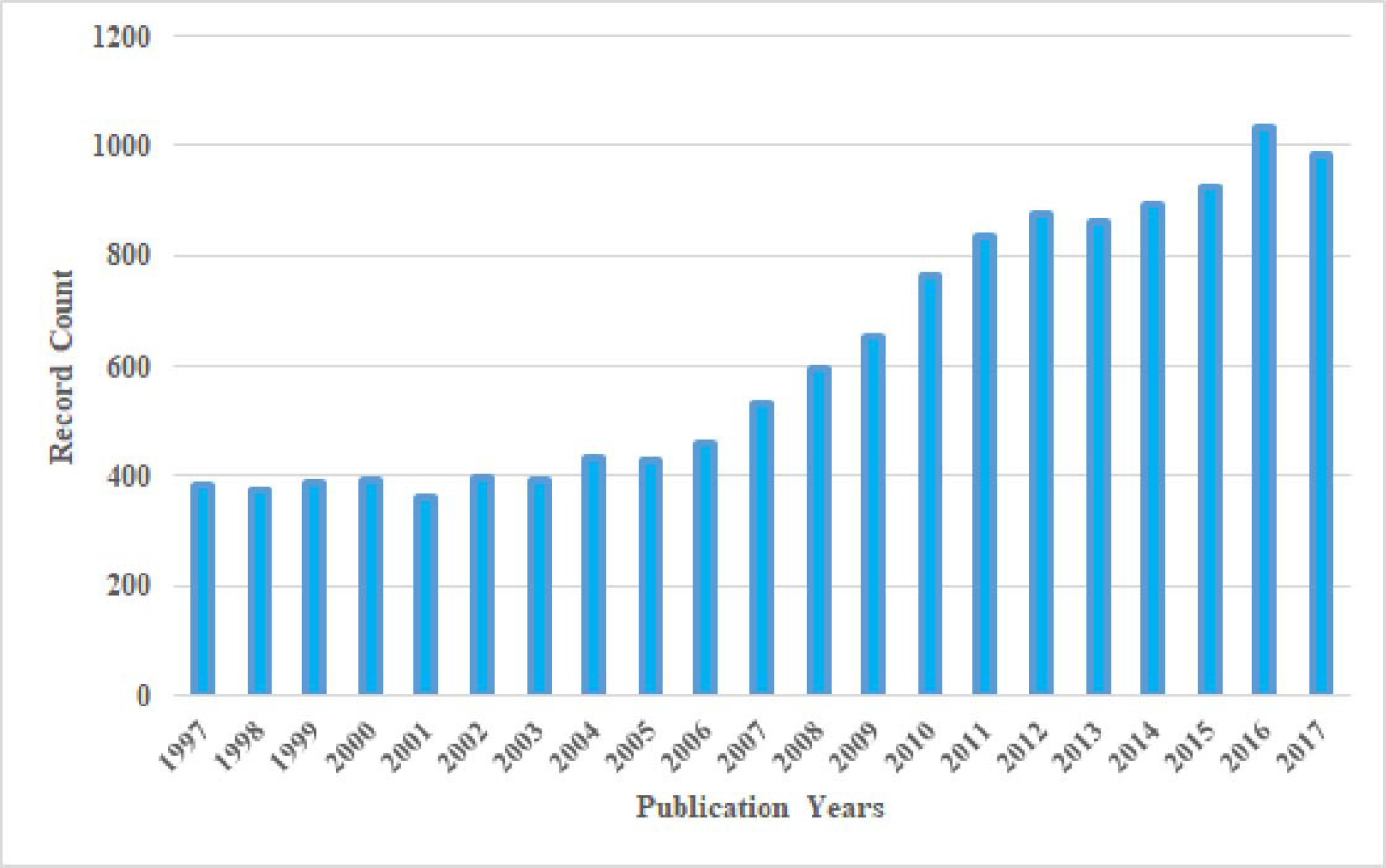
The number of publications related to *Trichoderma* in Wed of Science from 1997 to 2017

**Fig 2.**
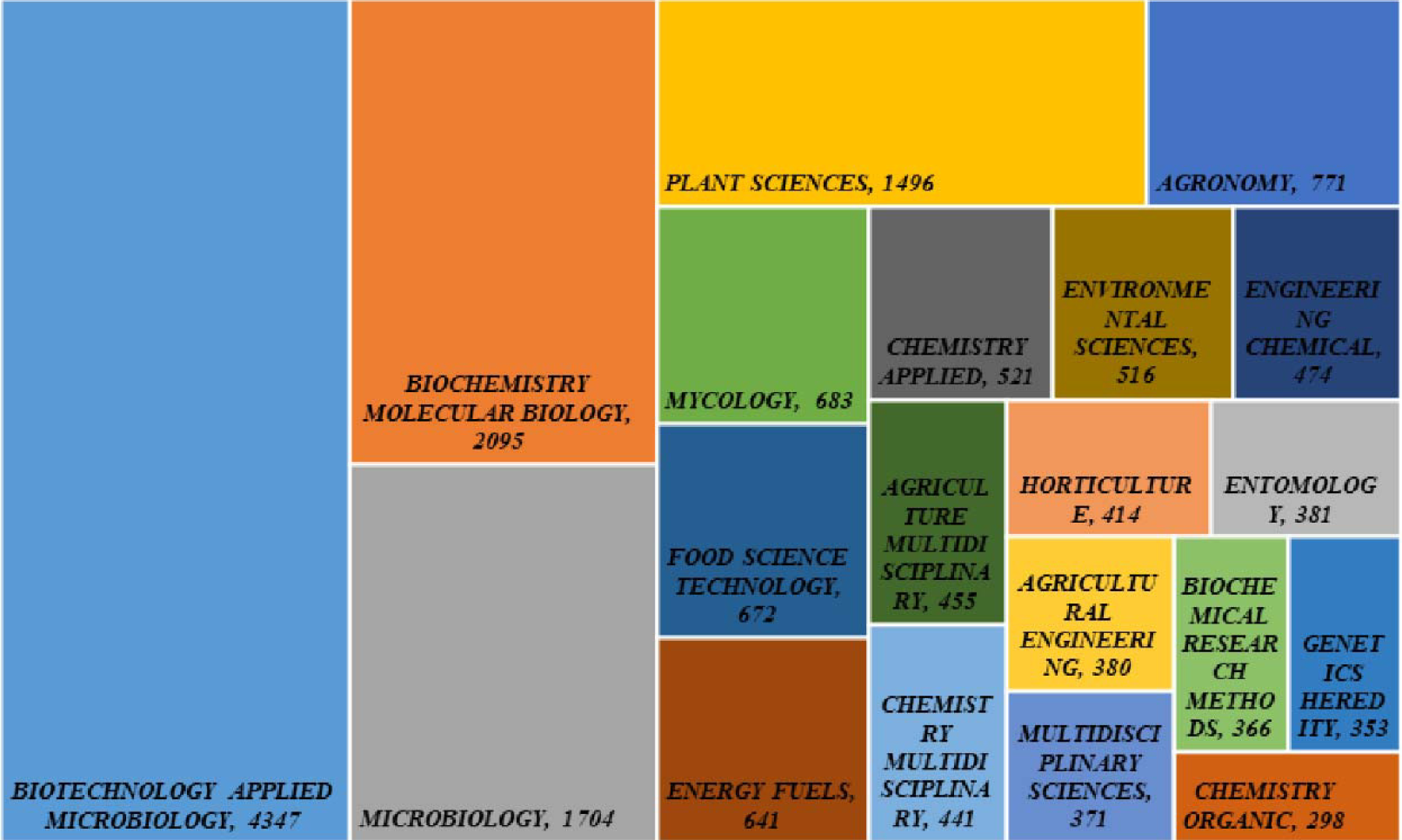
The research fields related to *Trichoderma* in Wed of Science from 1997 to 2017

In order to reduce the barrier standing between non-taxonomic researchers and *Trichoderma*-based researches, *Trichoderma* taxonomists have developed a series of identification tools. Samuels and colleagues collected and catalogued phenotypic characters of *Trichoderma* spp. to create an interactive key (http://nt.ars-grin.gov/taxadescriptions/keys/TrichodermaIndex.cfm, not available currently) for species identification. Although the interactive key makes comparison of different phenotypic characters convenient by avoiding referencing to protologue of each species, phenotype-based identification has inherent defects. For instance, the culture appearance of *Trichoderma* spp. is very susceptible to environmental factors without strict control, leading to difficulty to distinguish those phenotypic traits between species based on classical taxon system. The scenario result inevitably in the determination of phenotypic characters based on subjective judgement and homopasy characters of other species. What is more, the description of new *Trichoderma* spp. is rely heavily on DNA-based methods (Chen & Zhuang, 2017; Jaklitsch & Voglmayr, 2015a). These defects, not only exist in *Trichoderma* identification but commonly in fungi, make it difficult for a quick and accurate identification in species level even for experts.

To make some improvements, DNA barcoding, originally for species diagnosis in the animals (Hebert et al, 2003a; Hebert et al, 2003b), was introduced in *Trichoderma* identification by Druzhinina et al. and integrated with a program namely *Trich*Okey (Druzhinina et al, 2005). The original version of *Trich*OKey made use of several species-, clade- and genus-specific oligonucleotides sequences (named as hallmarks) derived from ITS (nuclear ribosomal internal transcribed spacer) region for a quick identification of 75 single species, 5 species pairs and 1 species triplet (Druzhinina et al, 2005). A more reliable version allowing simultaneous identification of multiple ITS sequences is in currently using (Druzhinina & Kopchinskiy, 2006) although the ITS region was currently considered to be insufficient for *Trichoderma* identification (Raja et al, 2017). Another identification tool box, *Tricho*BLAST, for *Trichoderma* identification had also been developed, which is a combination of a multilocus database of phylogenetic markers (MDPM), a diagnosis program of phylogenetic markers (*Tricho*MARK) and a local BLAST server (Kopchinskiy et al, 2005). *Trich*OKey and *Tricho*BLAST are *Trichoderma*-specific identification tools and had promoted *Trichoderma*-related studies (Matarese et al, 2012; Mohamed-Benkada et al, 2006) during an earlier period with the classification system in genus *Trichoderma* containing only 88 species (Kopchinskiy et al, 2005). However, the current taxonomy of *Trichoderma* has changed dramatically and a recent report listed 254 accepted *Trichoderma* names (Bissett et al, 2015) that have not been updated into the databases of *Trich*OKey and *Tricho*BLAST. Thus, there are no *Trichoderma*-specific identification tools currently applicable.

Apart from the mentioned identification databases, there are numerous databases with a broad diagnostic scope not only dedicate to *Trichoderma* (Thangadurai et al, 2016; Yahr et al, 2016). One of the most familiar databases is Genbank (share the same nucleotide database with DDBJ and EMBL). The Genbank database contains the largest number of nucleotide sequences including multilocus barcodes. However, identification of *Trichoderma* spp., similarly to other fungi, via BLASTn program in Genbank was advised to be cautious due to the non-curated association of sequences with species name (Druzhinina et al, 2005; Nilsson et al, 2006). Aware of this issue, a curated sequence database was created especially for *Trichoderma*, referred to as RefSeq Targeted Loci database (RTL), with a joint effort between the National Center for Biotechnology Information (NCBI) and fungal taxonomy experts (Robbertse et al, 2017). Besides, among the numerous curated databases involving fungi sequences, the UNITE (User-friendly Nordic ITS Ectomycorrhiza Database) (Urmas et al, 2005) could also be adopted to identify *Trichoderma* spp. within a limited number of species. For a comprehensive learning of the identification tools and databases, it is advised to refer to the latest publications (Raja et al, 2017; Thangadurai et al, 2016).

Taken together, the most suitable identification system currently available for *Trichoderma* is RTL-based BLASTn search due to its consistency with the up-to-date taxonomy of *Trichoderma*. Nevertheless, the RTL is mainly focused on the ITS region, which is not competent for identification at the species level of some speciose genera including *Trichoderma*. Moreover, there is no cross-referring tool at NCBI to make full use of its sequences resources efficiently for species identification. To overcome drawbacks of current taxonomic tools, a polyphasic identification system for *Trichoderma* (PIST) was constructed by utilizing a comprehensive information of nucleotide markers (ITS, TEF1 and RPB2) and phenotypic characters based on the latest *Trichoderma* taxonomy. PIST was run as an interactive system and is easily extended in database issue and even cover more fungi in other genus.

## Results

### Outline of the web interface

There are three blocks designed in the web interface currently available for users as shown in Fig 3.

**Fig 3.**
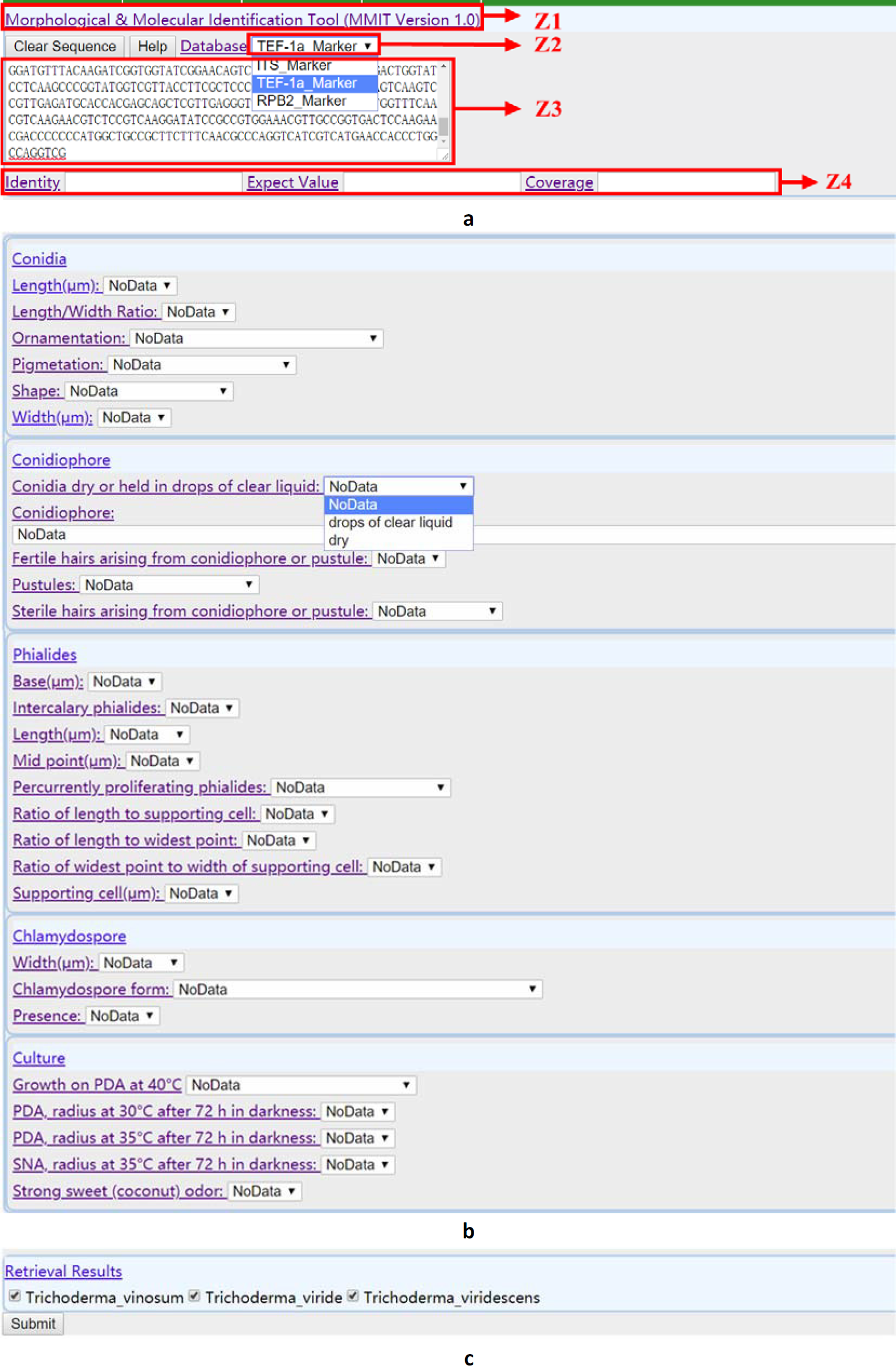
Outline of the web interface of PIST **a**, **b**, **c** exhibit Block 1, Block 2 and Block 3 of the interface separately. Z1, Z2 and Z3 in red color represent three operation zones in Block1, and the corresponding area was indicated in red boxes.

Block 1 (Fig 3a) contains four operating zones (Z1, Z2, Z3, Z4). Zone 1 is a short description of the identification system associated with a hyperlink, and function to initialize retrieval sets in the system by a single click. Zone 2 is a drop-down list of three nucleotide markers (ITS, TEF1, RPB2), most commonly used in *Trichoderma* identification, for users to choose a nucleotide marker database. Zone 3 is a textbox for users to input a nucleotide sequence corresponding to zone 2. The input sequence was required to only contain characters representing nucleic acids without any annotations and titles. Zone 4 includes three textboxes for users to change parameters of the BLAST program (Altschul et al, 1990). The parameter of Identity allowed an input of a number from 0 to 100 (i.e. 98) representing the sequence similarity from 0% to 100%. The parameter of Expect Value allowed an input of a number in Exponential notation (i.e. 1e-6), and the parameter of Coverage allowed an input of a number from 0.0 to 1.0 representing the sequence coverage from 0% to 100%. As there is no exact cutoff value of the parameter being universally competent for indicating conspecific taxa (O’Brien et al, 2005; Raja et al, 2017), the default parameters wfollowst up based on published articles as follow: 97% Identity, 80% Coverage and 1e-6 Expect Value (Nilsson et al, 2008; Raja et al, 2017).

Block 2 showed phenotypic characters of *Trichoderma*. It provided a specific content corresponding to a phenotypic character with a drop-down list for users to pick from.

Block 3 provided a submission function by a single click at the “submit” button. Subsequently, it listed candidate *Trichoderma* species that meet the users’ retrieval requirement.

### **Identified *Trichoderma* species through nucleotide markers**

In this study, 87 *Trichoderma* species were accurately identified with a specific combination of nucleotide markers (ITS, TEF1, RPB2) using default parameters (Table 1). Among the 86 *Trichoderma* species, 80 species were identified with one nucleotide marker. In detail, 66 species could be identified with TEF1, 46 species with RPB2 and 18 species with ITS. In comparison of the capability of the three nucleotide markers in differentiating species (excluding those species with partial markers), 19 *Trichoderma* species were only identified with TEF1 while 2 species were identified only by RPB2 or ITS. These results were in accordance with the published opinions, ITS region was not sufficient for species identification in some highly speciose genera including *Trichoderma* (Raja et al, 2017), and TEF1 has superior resolution than ITS especially in some species complex (Chaverri et al, 2015; Stielow et al, 2015).

**Table 1.**
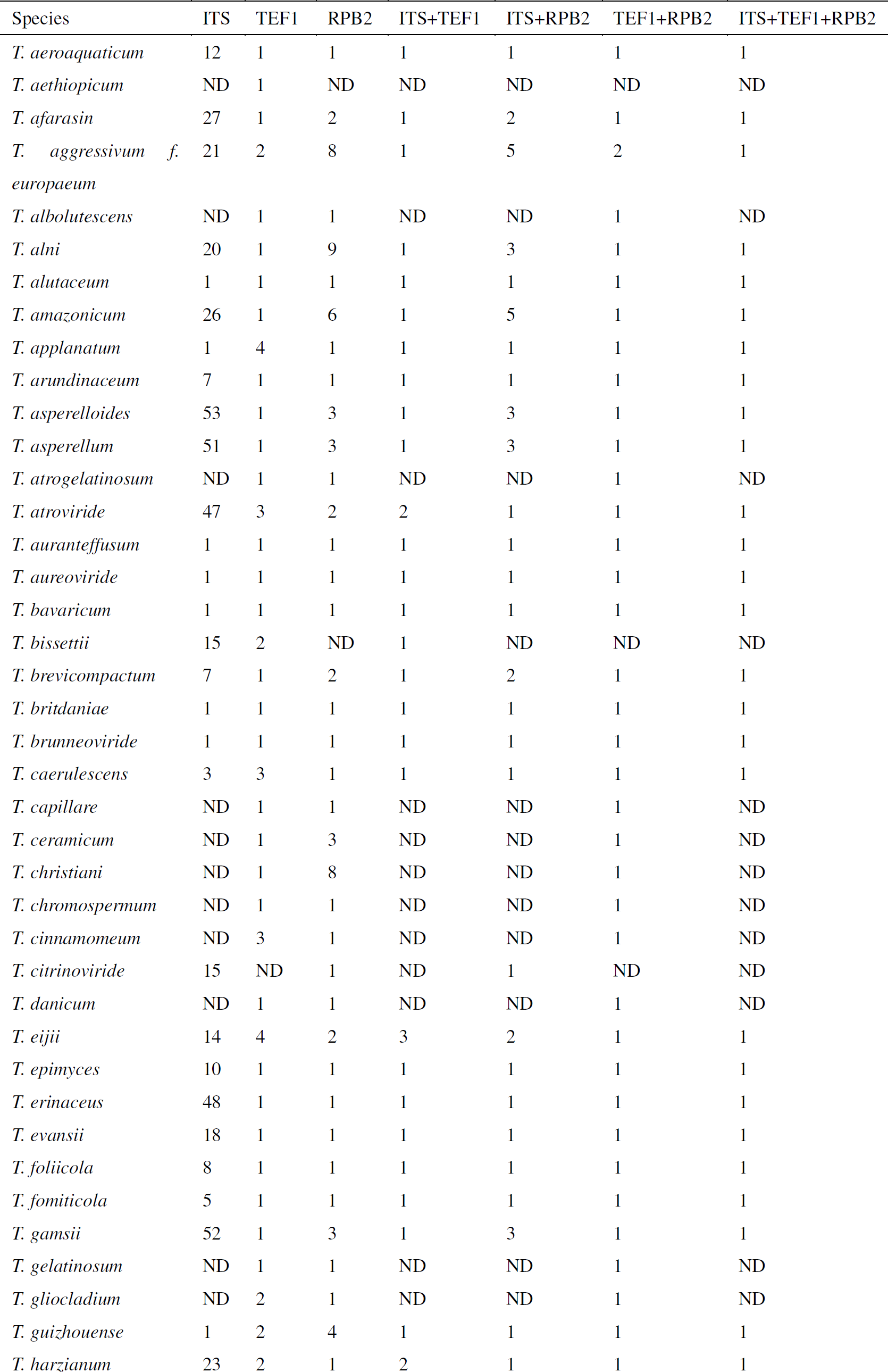
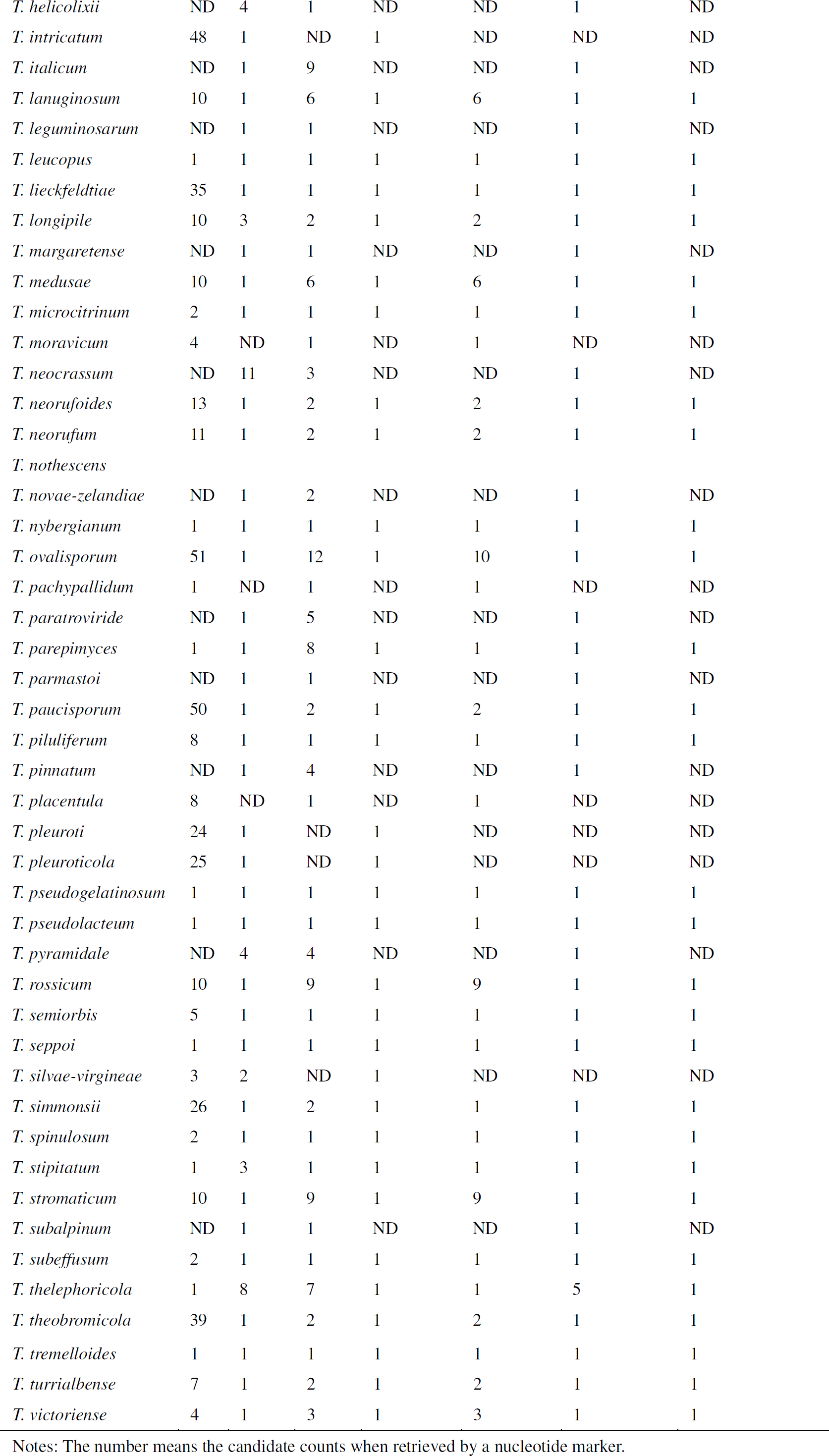
Accurately identified *Trichoderma* species through nucleotide markers with default retrieval parameters Notes: The number means the candidate counts when retrieved by a nucleotide marker.

For some *Trichoderma* species, the identification could only be completed by using a combination of different nucleotide markers. For instance, *T. aggressivum f. europaeum, T. bissettii* and *T. silvae-virgineae* were only identified by using the combination of ITS and TEF1, *T. eijii, T. neocrassum* and *T. pyramidale* by the combination of TEF1 and RPB2, *T. atroviride* and *T. longipile* by the combination of ITS and TEF1 or ITS and RPB2. These results reflected a supplementary effect of different nucleotide markers in *Trichoderma* identification.

### **Unidentified *Trichoderma* species through nucleotide markers**

There were 81 *Trichoderma* species with inseparable candidate species or no retrieval result if nucleotide markers with default parameters were used for retrieving (Table 2). For instance, no result comes out when *T. afroharzianum* was retrieved by using TEF1 unless changed the parameter of Coverage to 0.5. There are 2 candidates when retrieving *T. cremeum* even based on a combination of TEF1 and RPB2. After analyzing the corresponding marker sequences of the unidentified *Trichoderma* species, several types were divided as follows:

**Table 2.** Unidentified *Trichoderma* species through nucleotide markers with default retrieval parameters

| Species | ITS | TEF1 | RPB2 | ITS+TEF1 | ITS+RPB2 | TEF1+RPB2 | ITS+TEF1+RPB2 |
| --- | --- | --- | --- | --- | --- | --- | --- |
| *T. afroharzianum* | 27 | 1(0.5C) | 4 | 1 | 3 | 1 | 1 |
| *T. aggressivum* | 24 | 1(100I) | 1(100I) | 1 | 1 | 1 | 1 |
| *T. americanum* | 4 | 0 | 1(99I+0.6C) | 0 | 1 | 0 | 0 |
| *T. andinense* | ND | 1(96I) | 1(0.7C) | ND | ND | 1 | ND |
| *T. appalachense* | 51 | 1(99I) | ND | 1 | ND | ND | ND |
| *T. atlanticum* | 8 | 1(98I) | 5 | 1 | 3 | 1 | 1 |
| *T. atrobrunneum* | 27 | 1(0.5C) | 7 | 1 | 5 | 1 | 1 |
| *T. austriacum* | 1(100I) | 3 | ND | 1 | ND | ND | ND |
| *T. austrokonigii* | 48 | 1(93I) | 1(96I) | 1 | 1 | 1 | 1 |
| *T. camerunense* | 22 | 1(98I) | ND | 1 | ND | ND | ND |
| *T. caribbaeum* | 51 | 1(99I +0.3C) | 9 | 1 | 8 | 1 | 1 |
| *T. catoptron* | ND | 1(0.7C) | 2 | ND | ND | 1 | ND |
| *T. chlorosporum* | ND | 10 | 1(99I) | ND | ND | 1 | ND |
| *T. citrinum* | 8 | 0 | 4 | 0 | 4 | 0 | 0 |
| *T. compactum* | ND | 1(0.1C) | ND | ND | ND | ND | ND |
| *T. crassum* | 22 | 0 | 3 | 0 | 2 | 0 | 0 |
| *T. cremeoides* | ND | 2(99I) | 4(99I) | ND | ND | 1 | ND |
| *T. cremeum* | ND | 3(98I) | 3(99I) | ND | ND | 2 | ND |
| *T. dingleyae* | 49 | 1(0.4C) | 9 | 1 | 8 | 1 | 1 |
| *T. dorotheae* | ND | 1(98I) | 9 | ND | ND | 1 | ND |
| *T. effusum* | 0 | 0 | ND | 0 | ND | ND | ND |
| *T. estonicum* | 3 | 11 | 1(99I) | 2 | 1 | 1 | 1 |
| *T. eucorticoides* | 1 | 0 | 1 | 0 | 1 | 0 | 0 |
| *T. europaeum* | ND | 1(98I) | 4 | ND | ND | 1 | ND |
| *T. fertile* | ND | 1(98I) | 2 | ND | ND | 1 | ND |
| *T. flaviconidium* | 1(99I+0.3C) | 1 | ND | 1 | ND | ND | ND |
| *T. floccosum* | ND | 7(96I) | 4 | ND | ND | 1 | ND |
| *T. ghanense* | ND | 1(0.5C) | 1 | ND | ND | 1 | ND |
| *T. hamatum* | 19 | 0 | 1(98I+0.3C) | 0 | 1 | 0 | 0 |
| *T. helicum* | ND | 1(86I) | 1(0.7C) | ND | ND | 1 | ND |
| *T. hispanicum* | 51 | 1(98I) | ND | 1 | ND | ND | ND |
| *T. istrianum* | ND | 1(99I) | 8 | ND | ND | 1 | ND |
| *T. koningiopsis* | 51 | 1(96I) | 2(0.7C) | 1 | 2 | 1 | 1 |
| *T. lacuwombatense* | ND | 1(0.3C) | 4(0.7C) | ND | ND | 1 | ND |
| *T. mediterraneum* | ND | 1(98I) | 5 | ND | ND | 1 | ND |
| *T. mienum* | 2 | 1(0.6C) | 1(0.5C) | 1 | 1 | 1 | 1 |
| *T. minutisporum* | ND | 1(98I) | 5 | ND | ND | 1 | ND |
| *T. oblongisporum* | ND | 0 | 1(98I) | ND | ND | 0 | ND |
| *T. olivascens* | ND | 1(98I) | 16 | ND | ND | 1 | ND |
| *T. oligosporum* | 1(91I) | 1(96I) | 1 | 1 | 1 | 1 | 1 |
| *T. orientale* | ND | 1(96I+0.4C) | 4 | ND | ND | 1 | ND |
| *T. parapiluliferum* | 8 | 1 | 0 | 1 | 0 | 0 | 0 |
| *T. parareesei* | ND | 1(0.5C) | 2(0.7C) | ND | ND | 1 | ND |
| *T. parestonicum* | 3 | 1(99I) | 3 | 1 | 3 | 1 | 1 |
| *T. peltatum* | 1 | 1(96I) | 1(0.7C) | 1 | 1 | 1 | 1 |
| *T. petersenii* | 51 | 9(96I+0.3C) | 7 | 1 | 7 | 1 | 1 |
| *T. pezizoides* | 0 | 0 | 0 | ND | ND | ND | ND |
| *T. phyllostachydis* | ND | 1(0.4C) | 1(0.7C) | ND | ND | 1 | ND |
| *T. polysporum* | ND | 1(96) | 2(96I) | ND | ND | 1 | ND |
| *T. priscilae* | ND | 1(100I) | 4(99I) | ND | ND | 1 | ND |
| *T. protopulvinatum* | ND | 1(98I) | 4 | ND | ND | 1 | ND |
| *T. pseudokoningii* | 14 | 1(0.6C) | 0 | 1 | 0 | 0 | 0 |
| *T. pseudonigrovirens* | ND | 0 | 1 | ND | ND | 0 | ND |
| *T. pseudostramineum* | 0 | 0 | 0 | 0 | 0 | 0 | 0 |
| *T. psychrophilum* | 2 | 1(0.4C) | 1(0.7C) | 1 | 1 | 1 | 1 |
| *T. pubescens* | 21 | 0 | 1 | 0 | 1 | 0 | 0 |
| *T. reesei* | 15 | 1(0.6C) | 2(0.2C) | 1 | 2 | 1 | 1 |
| *T. rifaii* | 10(99I) | 3 | ND | 2 | ND | ND | ND |
| *T. rodmanii* | 9 | 1(0.4C) | 1(0.6C) | 1 | 1 | 1 | 1 |
| *T. rogersonii* | 41 | 1(0.4C) | 1 | 1 | 1 | 1 | 1 |
| *T. rufobrunneum* | 1(100I) | 2 | 9 | 1 | 1 | 2 | 1 |
| *T. samuelsii* | 50 | 1(98I) | 18 | 1 | 16 | 1 | 1 |
| *T. saturnisporopsis* | ND | ND | 1(99I+0.6C) | ND | ND | ND | ND |
| *T. saturnisporum* | ND | 1(0.6C) | 1 | ND | ND | 1 | ND |
| *T. sempervirentis* | ND | 1(99I) | 18 | ND | ND | 1 | ND |
| *T. sinense* | ND | 1(0.5C) | 1(0.6C) | ND | ND | 1 | ND |
| *T. sinoluteum* | 1 | 1(96I) | 2 | 1 | 1 | 1 | 1 |
| *T. sinuosum* | 4 | 1(100I) | 1(99I) | 1 | 1 | 1 | 1 |
| *T. spirale* | 21 | 1(0.4C) | 1 | 1 | 1 | 1 | 1 |
| *T. stilbohypoxyli* | 37 | 1(96I+0.4C) | 1(0.7C) | 1 | 1 | 1 | 1 |
| *T. stramineum* | 20 | 0 | 1 | 0 | 1 | 0 | 0 |
| *T. strigosellum* | 49 | 1(95I) | 2 | 1 | 2 | 1 | 1 |
| *T. strigosum* | 46 | 1(0.6C) | 1(0.7C) | 1 | 1 | 1 | 1 |
| *T. trixiae* | 46 | 1(99I) | 18 | 1 | 16 | 1 | 1 |
| *T. vinosum* | 51 | 2(98I) | 1(99I) | 2 | 1 | 1 | 1 |
| *T. viridarium* | ND | 1(98I) | 20 | ND | ND | 1 | ND |
| *T. viride* | 40 | 0 | 1(98I) | 0 | 1 | 0 | 0 |
| *T. viridescens* | ND | 1(99I) | 19 | ND | ND | 1 | ND |
| *T. viridialbum* | ND | 1(99I) | 18 | ND | ND | 1 | ND |
| *T. virilente* | ND | 1(99I) | 11 | ND | ND | 1 | ND |
| *T. voglmayrii* | ND | 1 | 1(0.5C) | ND | ND | 1 | ND |

Note: 1(96I+0.4C) means there is 1 candidate when retrieved by changing the parameter to 96% Identify and 0.4 Coverage. The numbers in green color mean the corresponding sequences belong to type 1, testing sequences share regional overlap with the database sequences. The numbers in orange color mean the corresponding sequences belong to type 2, testing sequences share less regional overlap with the database sequences and are not component

#### (T1) Overlap region of sequences contain barcode gap

There were 93 sequences within this type. It was supposed that the failing of identification of *Trichoderma* species with sequences of this type due to the regional overlaps between marker sequences used for testing and database construction. Essentially, the regional overlap was caused by the different primer pairs employed to amplify the nucleotide markers as there were more than one universal primer pairs designed for each nucleotide marker (Druzhinina & Kubicek, 2005; Liu et al, 1999; Martin & Rygiewicz, 2005a). In this case, a successful identification could be achieved by reducing the threshold of Coverage if the overlap region contained a barcode gap. A practical case showed that an accurate identification of *T. afroharzianum* succeeded by setting the Coverage of TEF1 as 0.5, however, there was no retrieval result under the default parameters. Another failure of species identification is caused by the intraspecific variability of sequence similarity. Up to now, within *Trichoderma* species, there is no defined bound of sequence similarity for each nucleotide marker. On this condition, a successful identification could be accomplished by adjusting the threshold of Identity (i.e. increasing the value of Identity to 100 while retrieving *T. austriacum* with ITS or decreasing that to 96 while retrieving *T. andinense* with TEF1). In some cases, a simultaneous adjusting of Identity and Coverage was needed for final identification (i.e. setting coverage as 0.6 and Identity as 99 when using RPB2 for *T. americanum* identification).

#### (T2) Overlap region of sequences did not contain barcode gap

There are 18 sequences in this type and as much as 13 are TEF1 as it contains three regions (intron 4, intron 5 and exon 6) usually amplified by researchers (Druzhinina & Kubicek, 2005). A practical case showed that the TEF1 sequence for PIST testing from *T. citrinum* was located at intron 4 and 5, while the corresponding sequence deposited in database located at exon 6. Consequently, the two sequences were incomparable and was unable to be employed for identification of *T. citrinum*. Except for TEF1 marker, there were four RPB2 sequences from *T. parapiluliferum, T. pezizoides, T. pseudokoningii* and *T. pseudostramineum*, and one ITS sequence from *T. pezizoides* shared less overlap regions with the corresponding sequences in database and were not enough for species identification.

#### (T3) Tested marker sequences showed low sequence similarity intraspecies

There were two tested marker sequences in this type and both are ITS markers sourced from *T. effusum* and *T. pseudostramineum* separately (Fig 4). Those sequences showed an excessively low similarity with corresponding sequences deposited in databases. As all the sequences were from type material or submitted by *Trichoderma* taxonomists, the lower similarity of compared sequences was unlikely caused by the error taxonomic annotation. For ITS from *T. effusum*, there are a string of undetected nucleotides among the sequence (Fig 4a). That may be caused by the sequencing errors. For ITS from *T. pseudostramineum*, there are much INDEL mutation between the two sequences used for species identification (Fig 4b). The reality of the mutation needs to be confirmed from the corresponding strains.

**Fig 4.**
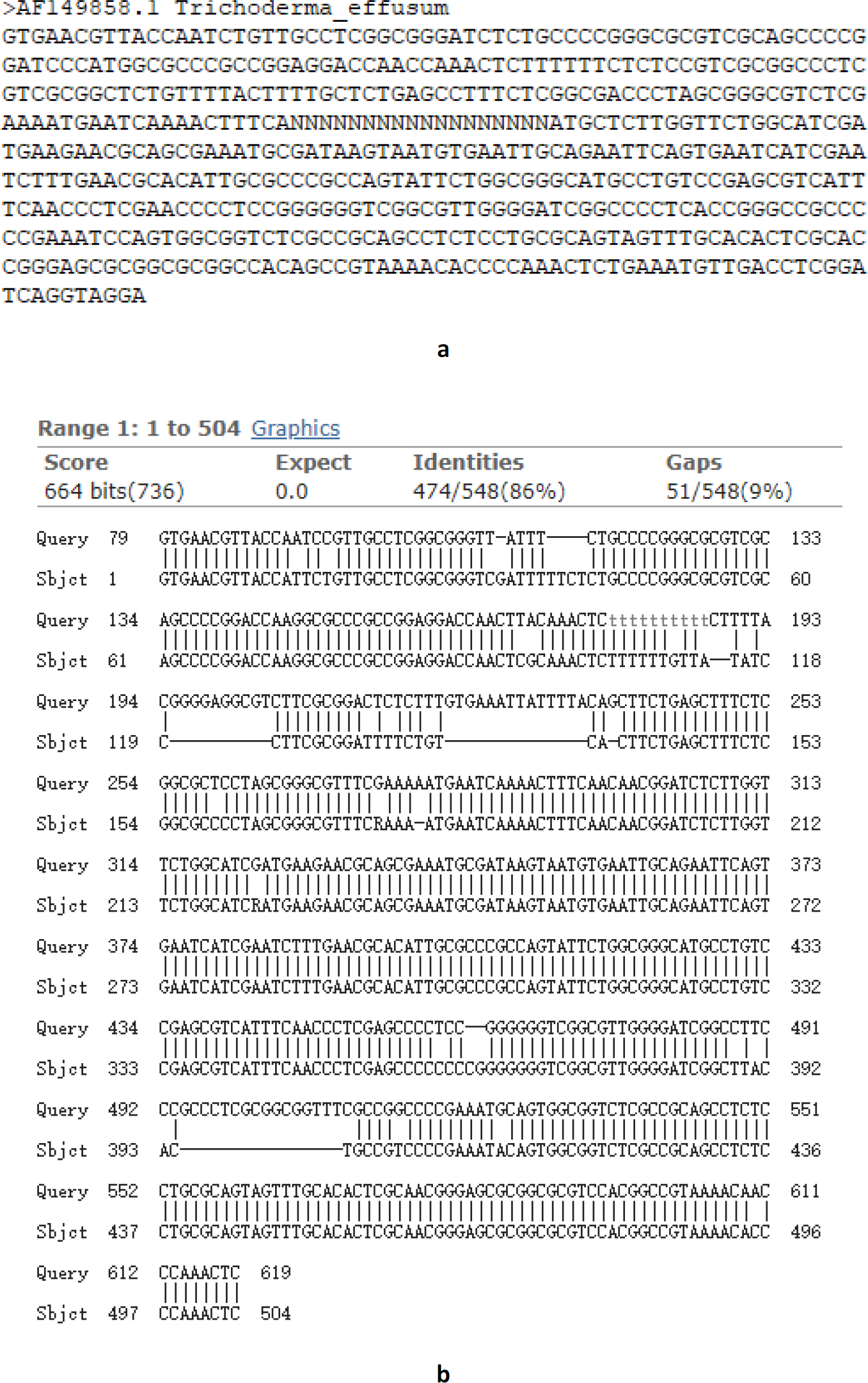
Tested marker sequences showed low sequence similarity intraspecies **a** exhibits the tested ITS sequence containing undetected nucleotides from *T. effusum* **b** exhibits the sequence alignment of ITS sequences from two *T. pseudostramineum* strains. Query sequence is the tested ITS from *T. pseudostramineum* TUFC 60104 with an accession number of ND134435. Sbjct sequence is ITS deposited in nucleotide marker databases from *T. pseudostramineum* GJS 90-74 with an accession number of DQ835420.

### **Misidentified *Trichoderma* species through nucleotide markers**

Compared with the unidentified *Trichoderma* species, the misidentified *Trichoderma* species have to be made for further analysis in detail. There were 20 *Trichoderma* species misidentified through one or more tested marker sequences as shown in Table 3. After analyzing the sequence similarity and genome locus of the marker sequences used for testing and database construction, the reason of the misidentification was supposed to be the lacking of barcode gaps in overlap region. And meanwhile, the overlap region exhibits a higher sequence similarity interspecies than intraspecies. For instance, testing of TEF1 sequences sourced from *T. ceraceum, T. corneum, T. deliquescens, T. lixii, T. longibrachiatum, T. pezizoides, T. pulvinatum, T. strictipile, T. sulphureum, T. tawa, T. tomentosum* and *T. virens*, RPB2 sequences from *T. cerinum, T. composticola, T. corneum, T. koningii, T. neosinense, T. paraviridescens, T. pulvinatum, T. crystalligenum*, and *T. strictipile*, and ITS from *T. crystalligenum* and *T. neotropicale*. The tested TEF1 from *T. barbatum* and RPB2 from *T. lentiforme* did not contain barcode gaps for species identification even though they share same genome loci with the corresponding sequences in nucleotide database.

**Table 3.** Misidentified *Trichoderma* species through nucleotide markers with default retrieval parameters

| Species | ITS | TEF1 | RPB2 | ITS+TEF1 | ITS+RPB2 | TEF1+RPB2 | ITS+TEF1+RPB2 |
| --- | --- | --- | --- | --- | --- | --- | --- |
| *T. barbatum* | 10 | <u>1</u> | 1(99I) | <u>1</u> | 1 | 0 | 0 |
| *T. ceraceum* | ND | <u>3</u> | 1(98I) | ND | ND | 0 | ND |
| *T. cerinum* | ND | 1(99I+0.3C) | <u>2</u> | ND | ND | 0 | ND |
| *T. composticola* | 47 | 1(0.4C) | <u>5</u> | 1 | <u>5</u> | 0 | 0 |
| *T. corneum* | ND | <u>1</u> | <u>11</u> | ND | ND | <u>1</u> | ND |
| *T. crystalligenum* | <u>1</u> | 0 | <u>1</u> | 0 | <u>1</u> | 0 | 0 |
| *T. deliquescens* | 4 | <u>1</u> | 2 | <u>1</u> | 2 | <u>1</u> | <u>1</u> |
| *T. koningii* | 49 | 1 | <u>1</u> | 1 | <u>1</u> | 0 | 0 |
| *T. lentiforme* | 24 | 1(96I) | <u>3</u> | 1 | <u>2</u> | 0 | 0 |
| *T. lixii* | 24 | <u>2</u> | ND | 0 | ND | ND | ND |
| *T. longibrachiatum* | 15 | <u>1</u> | 3 | 0 | 1 | <u>1</u> | 0 |
| *T. neosinense* | 48 | 0 | <u>4</u> | 0 | <u>4</u> | 0 | <u>2</u> |
| *T. neotropale* | <u>22</u> | 1(90I) | ND | 0 | ND | ND | ND |
| *T. paraviridescens* | 46 | 1(0.4C) | <u>4</u> | 1 | <u>4</u> | 0 | 0 |
| *T. pulvinatum* | 4 | <u>1</u> | <u>3</u> | <u>1</u> | <u>3</u> | <u>1</u> | <u>1</u> |
| *T. strictipile* | ND | <u>8</u> | <u>1</u> | ND | ND | 0 | ND |
| *T. sulphureum* | ND | <u>1</u> | 1(99I) | ND | ND | 0 | ND |
| *T. tawa* | ND | 0 | <u>1</u> | ND | ND | 0 | ND |
| *T. tomentosum* | 24 | <u>4</u> | 1(98I) | <u>2</u> | 1 | <u>1</u> | <u>1</u> |
| *T. virens* | 21 | <u>9</u> | 1(99I) | <u>1</u> | 1 | <u>1</u> | <u>0</u> |

Note: The numbers in underling and bold mean the retrieval candidates do not contain the right *Trichoderma* species. Others notes are the same with Table 1.

Considering the misidentified cases, it is suggested that with a specific retrieved candidate species, more characters should be employed to confirm the identification. A reliable species identification in PIST was based on the consistent identification of all nucleotide markers and phenotypic characters.

## Discussion

The purpose of this study is to develop a polyphasic identification system for genus *Trichoderma* to keep in step with the burgeoning taxonomy system and facilitate *Trichoderma*-based research. Consequently, serious databases covering 252 *Trichoderma* species were constructed, including 338 ITS sequences, 435 TEF1 sequences, 415 RPB2 sequences and 28 phenotypic characters. In addition, this is the first report to adopt a polyphasic identification system for *Trichoderma* species identification. Since being introduced by Persoon in 1794 (Druzhinina & Kubicek, 2005; Persoon, 1794), *Trichoderma* taxonomy and species identification are difficult issues. The abundant homoplasy in phenetic characters is likely the reason (Druzhinina et al, 2006). Therefore, the taxonomy and identification of *Trichoderma* have evolved from a single phenotypic character-based method to a molecular character-based method supplemented by phenotypic observation. Along the conversion, the major challenges have to be conquered, including unstable multiple phenotypic features and insufficient DNA barcode information. As the rapid changing in *Trichoderma* taxonomy, the species counts are continuously increasing and the current identification systems are not suitable for an accurate and efficient identification (Robbertse et al, 2017). Referring to the identification systems of speciose genera *Penicillium* and *Aspergillus* (http://www.westerdijkinstitute.nl/Aspergillus/Biolomicsid.aspx), the poly-phasic-based species identification is a promising method for *Trichoderma*.

### Comparison of different identification methods

Phenotypic characters are traditionally employed to form the fungi taxonomic system and to identify fungi species in the manner of a phenotype-based species key. It is well documented that phenotypic characters is not efficient and effective as molecular characters in species or genus identification (François et al, 2004; Raja et al, 2017). For genus *Trichoderma*, the resolution of phenotypic characters in species identification is just limited within 27 species (Bissett, 1984; Bissett, 1991a; b; c; 1992). However, this does not mean that the phenotypic method could be replaced by molecular method for species identification. The irreplaceability could be exemplified by the identification process of *T. cremeum* (Table 2) through PIST. Just by retrieving with a combination of TEF1 (98% Identity) and RPB2 (99% Identity) sequences, *T. cremeum* was not separated from *T. surrotundum*, but the good distinction of both species could be completed if a character of conidia shape (subglobose to ovoidal) was involved in the identification. This is why nowadays the integral of phenotypic and molecular characters was still needed to confirm a new *Trichoderma* species (Jaklitsch, 2011; Jaklitsch & Voglmayr, 2015b; Zhang et al, 2018b).

The DNA barcode is an effective tool for species identification initiated from the animal kingdom and then be introduced into fungi. To meet the requirement of a quick and accurate identification of *Trichoderma* species, ITS region-based barcode method was developed, named as *Trich*OKey, which is able to operate identification of 75 individual species (Druzhinina et al, 2005). However, the system is not powerful enough as more new *Trichoderma* species emerged. The ITS region was employed for fungal identification in almost all fungal databases (Geiser et al, 2004; Robert et al, 2011; Yahr et al, 2016) and nominated as the official fungal barcode (Schoch et al, 2012). However, as the number of species continuously increasing in genus *Trichoderma*, 100 species in year 2006 and 256 species in 2015 (Bissett et al, 2015; Druzhinina et al, 2006), the ITS region could not bear a heavier load of species identification due to its natural resolution. A concept of second barcode was proposed to deal with the speciose genera, including *Trichoderma* (Samson et al, 2014; Stielow et al, 2015). Although the *Tricho*BLAST program (Kopchinskiy et al, 2005) contains multiloci markers for distinction of *Trichoderma* species, the retrieval processes of each marker are separated and lacking a cross-referring procedure among different marker databases, and the same problem also exists in Genbank (Robbertse et al, 2017). A secondary metabolite profile-based chemotaxonomy method was also employed in *Trichoderma* identification, but with a small scale of only 7 species (Kang et al, 2011). Thus currently or in a shortcoming days the characters cannot be applied in the taxonomy of *Trichoderma* species due to the overcomplexity of secondary metabolites among species.

Despite of the phylogenetic tree-based identification, superior in defining new species, but not convenient for identifying existed species, the polyphasic identification method has an advantage of comprehensive utilization of different characters (i.e. phenotype and molecular) particularly in combination with computer assistant system. A series of polyphasic identification systems have been developed for *Aspergillus, Penicillium, Fusarium* (http://www.westerdijkinstitute.nl/Collections/), *Ceratocystis, Colletotrichum* and *Phytophthora* (http://www.q-bank.eu/fungi/). All the polyphasic identification systems mentioned above are derived from BioloMICS software (Robert et al, 2011) and allow a simultaneous submission of several different characters. And yet, no polyphasic identification system was designed for genus *Trichoderma*.

PIST is expected to be a polyphasic system for *Trichoderma* identification. The core part of PIST is a program coded in Perl language, making the cross referring of different kinds of databases practicable. Unlike other polyphasic identification methods, the retrieval of characters in PIST is achieved in a step-by-step way. By this means, users are able to analysis the retrieval results after every submission action, and this is helpful especially deal with different nucleotide markers. During the testing process of PIST, different nucleotide markers and phenotypic characters showed complementary functions in species identification and turned out to identify 175 *Trichoderma* species, indicating the advantage of the polyphasic method. There are only 13 species (*T. citrinum, T. corneum, T. crassum, T. crystalligenum, T. deliquescens, T. effusum, T. lixii, T. neosinense, T. pezizoides, T. pseudostramineum, T. pulvinatum, T. strictipile, T. tawa*) cannot be identified due to lacking of enough sequences information.

### Problems in the identification process using molecular characters

Although the molecular character-based identification of fungi in species level is popular and predominant in taxonomy, there are problems needed to be resolved. One problem was the confounding of unverified and unqualified marker sequences deposited in open databases. This problem has attracted attention of taxonomist for a long time (D. et al, 2003; Druzhinina et al, 2005; Nilsson et al, 2006; Rytas, 2003). Fortunately, the problem has been successfully resolved since the numerous curated databases containing kinds of fungi nucleotide markers have been generated.

The current obstacle in species identification is mainly caused by the inconsistency of the amplified fragments from nucleotide markers, and results in an uneven sequence alignment. Consequently, the overlap regions between query sequences and subject sequences are likely to lose the barcode gaps or limited by retrieval parameters which in the end leading to a failure of species identification (Table 2), or even worse, an imperceptible misidentification (Table 3). To solve this problem, it was suggested that a unity of primer pairs used for amplifying nucleotide markers with the purpose of species identification. For genus *Trichoderma*, primer pairs of ITS4 and ITS5 for ITS marker, EF1728F and TEF1LLErev for TEF1 marker, fRPB2-5f and fRPB2-7cr for RPB2 marker are competent for species identification as they were able to generate amplicons with good resolution. (Jaklitsch, 2011; Jaklitsch & Voglmayr, 2015b; Martin & Rygiewicz, 2005b; Schoch et al, 2012).

### Prospective

A useful identification system depends heavily on qualified character databases. Currently, in the public databases, although the error characters are being corrected, a continuous update of databases is essential. For PIST, a complete set of phenotypic characters covering all species and standardized nucleotide databases will further improve the efficiency and accuracy of species identification. To achieve this goal, the assistance from *Trichoderma* taxonomist is imperative.

As the ever-growing requirement of identification even below the species level, such as an aggregation of *Trichoderma* strains with superior chitinase formation (Nagy et al, 2007), more nucleotide markers or characters from new dimensions will be developed and employed for identification. The polyphasic method will show a significant advantage and broad applications.

## Materials and methods

### Nucleotide marker databases

There are 213 ITS, 252 TEF1, and 243 RPB2 marker sequences representing 252 *Trichoderma* species included in the initial nucleotide marker databases (Appendix Table 1). All the marker sequences were picked out from the Genbank database mainly referring to a recently published article where a list of accepted *Trichoderma* names, reference strains and marker sequence accession numbers were proposed by *Trichoderma* taxonomists (Bissett et al, 2015). Actually, some marker sequences (Appendix Table 2&3) published in the reference article are obviously ambiguous for taxonomy use (Bissett et al, 2015). In these cases, marker sequences submitted from culture collections by *Trichoderma* taxonomists (i.e. John Bissett, Walter Gams, Walter Jaklitsch, Gary J. Samuels, Christian Kubicek, WY Zhuang) or certificated from type materials by NCBI were chosen as replacements.

### Phenotype database

The phenotypic characters used for database construction are partially adopted from the interactive key created by Samuels and colleagues. This database contains 28 characters from 46 *Trichoderma* species, and stored in a text file. All the *Trichoderma* names were in accordance with the currently accepted names (Bissett et al, 2015).

### Identification program

The PIST system is running in an Ubuntu Linux 16.04 server maintained by Center for Culture Collection of *Trichoderma* in Shanghai Jiao Tong University, and available through a web interface (http://mmit.china-cctc.org/). The operating environment is supported by Apache2, PHP5.3.3, BLAST program (Altschul et al, 1990), Perl and Bioperl. The interactive function between users and the web interface is achieved using PHP language. Nucleotide marker databases were formatted by BLAST program (Altschul et al, 1990) and stored separately according to the markers. The core cross-referring program was written in Perl language with Bioperl package, and perform the following functions:

1. Retrieving corresponding databases (in the manner of calling BLAST program for nucleotide marker databases, and field matching for phenotypic database) according to the submission of users and return species names as a result.
2. Adjusting all databases in limited scopes in accordance with the latest retrieval results.
3. Preparing the limited databases for the next retrieve unless initialization.
4. Adjusting the parameters (i.e. identity, coverage) when retrieving nucleotide marker databases by BLAST program (Altschul et al, 1990).
5. Initializing for a new retrieval.

### Verification of PIST

Totally 125 ITS, 183 TEF1, 172 RPB2 marker sequences from 188 *Trichoderma* species were used for determining the performance of PIST (Appendix Table 4) in consideration of the limitation of corresponding marker sequences available. The principles for selecting marker sequences and *Trichoderma* strains are the same with those for selecting marker sequences in nucleotide marker databases. In addition, for one *Trichoderma* species, the three markers should be of the same strain in order to determine the accuracy of PIST. The loci of the marker sequences were determined by *Tricho*MARK (Kopchinskiy et al, 2005). All kinds of possible combinations of the three nucleotide markers were used for *Trichoderma* spp. identification. After the verification process, all the tested sequences were integrated into the corresponding nucleotide marker databases.

## Acknowledgement

This work was supported by The National Key Research and Development Program of China (2017YFD0201108, 2017YFD0200403); Sino-US innovation program in key technology of crop disease and pest biological control; “948”key project of the Ministry of Agriculture of China (2016-X48); China Agricultural Research System (CARS-02); Intergovernmental Key International Scientific and Technological Innovation Cooperation (2017YFE0104900). Dr. He Meng and Weixing Ye are thanked for the support of software development.

